# Patient-derived models of brain metastases recapitulate the histopathology and biology of human metastatic cancers

**DOI:** 10.1101/2020.11.26.400036

**Authors:** Claudia C. Faria, Carlos Custódia, Rita Cascão, Eunice Paisana, Tânia Carvalho, Pedro Pereira, Rafael Roque, José Pimentel, José Miguéns, João T. Barata

## Abstract

**Purpose:** Dissemination of cancer cells from primary tumors to the brain is observed in the great majority of cancer patients, contributing to increased morbidity and being the main cause of death. Most mechanistic and preclinical studies have relied on aggressive cancer cell lines, which fail to represent tumor heterogeneity and are unsuitable to validate therapies due to fast cancer progression *in vivo*.

**Experimental design:** We established a unique library of subcutaneous and intracardiac patient-derived xenografts (PDXs) of brain metastases (BMs) from eight distinct primary tumor origins. Cancer progression in mice was compared to the matched patient clinical outcome, metastatic dissemination pattern and histopathological features. Preclinical studies with FDA approved drugs were performed.

**Results:** *In vivo* tumor formation of flank-implanted BMs correlated with patients’ poor survival and serial passaging increased tumor aggressiveness. Subcutaneous xenografts originated spontaneous metastases in 61% of the cases, including in the leptomeningeal space (21%). The intracardiac model increased the tropism to the brain and leptomeninges (46%). Strikingly, 62% of intracardiac PDXs shared metastatic sites with the donor patients, including the primary cancer organ and the central nervous system (CNS). Of therapeutic relevance, PDX-derived cultures and corresponding mouse xenografts can be effectively treated with targeted anticancer drugs.

**Conclusions:** Patient-derived models of BMs recapitulate the biology of human metastatic disease and can be a valuable translational platform for precision medicine.

**TRANSLATIONAL RELEVANCE:** Subcutaneous and intracardiac mouse xenografts of human brain metastases exhibit a spontaneous dissemination pattern that resembles patients’ metastatic disease. The preclinical testing of targeted anticancer drugs using patient-derived cultures and patient-derived xenografts of brain metastasis showed an effective therapeutic response. These translational models represent an outstanding tool to advance the understanding of the biology of brain metastases and to foster the rapid discovery of novel therapeutics.

## INTRODUCTION

Metastases are the main cause of cancer morbidity and mortality. Metastatic cancer cells can arise within the brain parenchyma (~40% of cancer patients)[1, 2] or spread to the leptomeninges to coat the brain and the spinal cord (~5–8% of cancer patients)[3, 4]. Both clinical conditions are associated with a dismal outcome with a median survival rate of 4–6 weeks for untreated patients and 8-16 weeks after standard of care therapies (palliative radiotherapy and chemotherapy)[5–7]. The primary tumors with the highest predisposition to develop brain metastases (BMs) include lung (40–60%), breast (15–30%) and melanoma (5–15%)[2, 8–11]. Despite the recent medical advances in the treatment of primary tumors, the incidence of fatal BMs has increased. This augmented incidence is most likely due to improved patient survival and thus time for dissemination, inefficacy of primary tumor therapies in treating BMs[12, 13], lack of available drugs that penetrate the blood–brain barrier (BBB)[14–17], and to the selection of chemoresistant metastatic clones upon treatment with DNA damaging agents[14, 15, 18, 19].

Although specific genes have been associated with BMs[20-24], the biological processes underlying the dissemination of cancer cells into the brain are not fully understood and there is a tremendous unmet medical need for the development of targeted therapies. One of the main caveats in advancing the treatment of BMs is the lack of appropriate biological models to study the disease. Most published studies have used cancer cell lines *in vitro*, minimizing tumor heterogeneity and not considering the complex host/cell interactions. Additionally, conventional preclinical cancer models, such as xenografts with immortalized cell lines, do not fully resemble the human tumor biology and thus have limited predictive power[25-29]. Genetic mouse models also fail to recapitulate the heterogeneity of primary human cancers and are limited by complex breeding schemes, incomplete tumor penetrance and variable tumor onset[30]. The recent generation of patient-derived xenografts (PDXs), by transplantation of patient tumor fragments subcutaneously or orthotopically into immune-deficient mice[31], emerged as a promising tool to better model human disease. These models recapitulate the main histological and genomic features of the parental tumors[25, 27, 32] at a relatively low passage[33], increasingly attracting interest to study disease mechanisms and therapeutic resistance[25, 27, 34]. They have also been used to identify and screen new drugs[32], and as a platform for personalized medicine, allowing patient’s tumor characterization and treatment selection. The ability of PDXs to maintain the phenotypic and molecular signature of human tumors has been described for different types of cancer, namely neuroblastoma[30, 35], glioblastoma[36], gastric cancer[37], breast cancer[38-40], lung cancer[41-43], pancreatic ductal adenocarcinoma[44-46], colon cancer[47, 48], melanoma[49], ovarian carcinoma[50, 51] and renal cancer[52, 53]. However, few publications have reported data from PDXs developed upon implantation of human BMs, which were derived from only a limited spectrum of cancer types, namely non–small cell lung cancer (NSCLC)[41] and breast cancer[54-56].

Taking advantage of the privileged access to BM samples of patients with detailed clinical annotation, we have successfully developed a unique library of PDXs of BMs from eight distinct primary tumor origins. We have developed subcutaneous and intracardiac models, performed a histopathological characterization of the tumors and thoroughly evaluated the pattern of cancer cell dissemination with particular emphasis in the central nervous system (CNS). To further validate our patient-derived models of BMs as a tool for precision medicine, we demonstrated the efficacy of FDA approved drugs in treating patient-derived cultures and the correspondent mouse xenografts. To our knowledge, this is the first study describing a spontaneous metastatic phenotype in PDXs of BMs, faithfully recapitulating human disease, and with an effective response to targeted anticancer therapies. We believe our patient-derived models of BMs constitute a clinically relevant platform for preclinical drug discovery and a powerful tool for personalized medicine.

## RESULTS

### Patient-derived tumor models reflect the clinical spectrum of human brain metastatic disease

Thirty-four surgical specimens were consecutively collected from patients with metastatic disease in the brain, skull and orbit, from ten different primary cancers (**Table 1**), in the Department of Neurosurgery at Hospital de Santa Maria (CHULN, Lisbon, Portugal). Tumor fragments from each specimen were implanted in the flank of NSG mice within 1 hour after surgery. Subcutaneous tumors were serially passaged *in vivo* until passage four and, in parallel, cancer cells were dissociated from tumor specimens to generate PDCs of BMs and intracardiac PDXs upon injection of cancer cells into the heart (left ventricle) of mice (**Figure 1A**). Upon reaching the humane endpoint, animals were euthanized, and all organs were collected for histopathological analysis and to evaluate the presence of metastases. PDCs were successfully generated from 61% of the engrafted tumors (**Table 1**).

**Figure 1.**
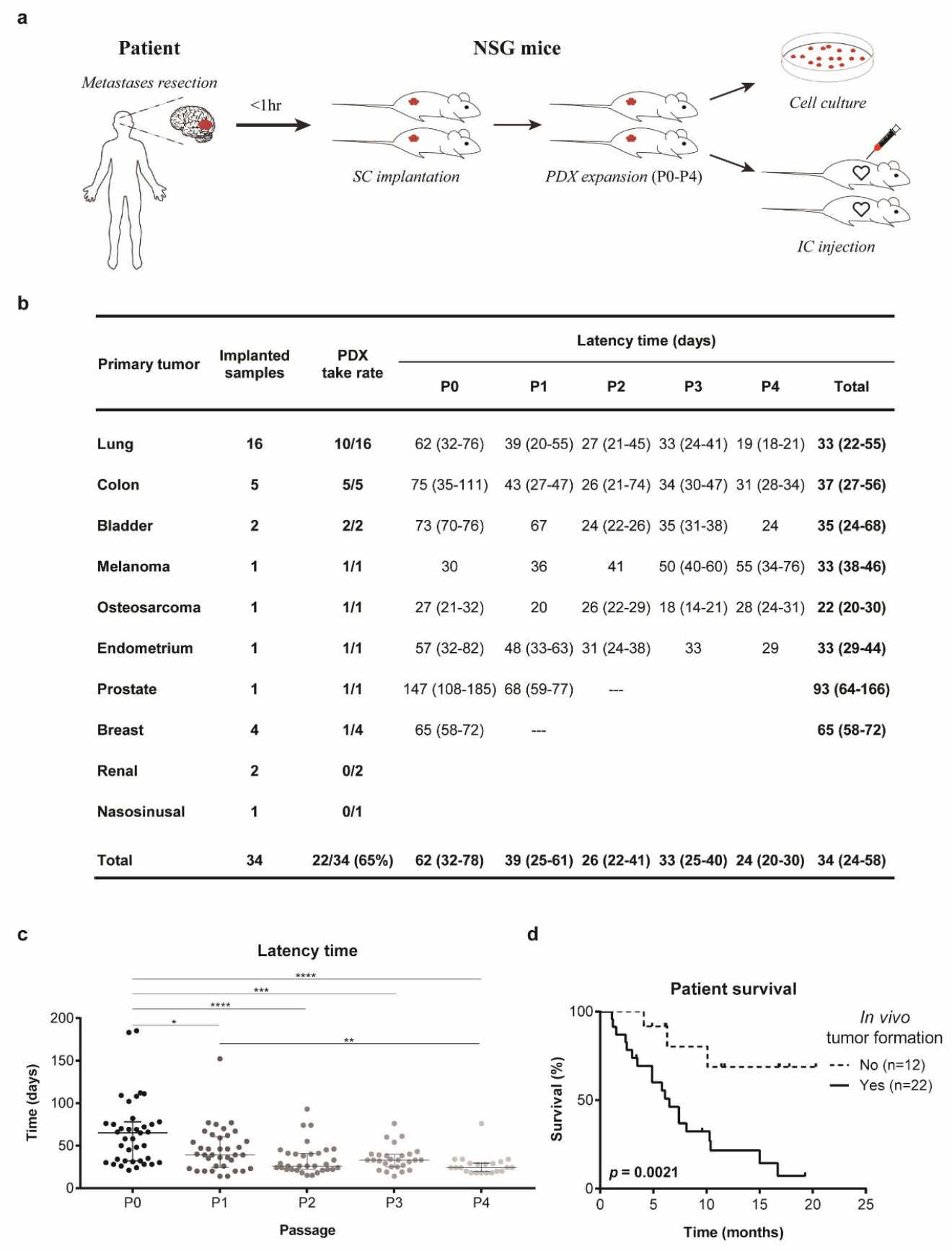
Subcutaneous xenografts derived from BMs surgical samples. (**A**) Experimental workflow for patient-derived models. Samples were implanted in the flank of immuno-compromised NSG mice and serially expanded until passage four. Flank tumors were dissociated into single cell suspensions that were primarily cultured or injected in the left cardiac ventricle of NSG mice. (**B**) Take rate and latency time of subcutaneously implanted human BMs from diverse primary cancers. (**C**) Tumor latency time decreases upon *in vivo* serial passaging. (**D**) The *in vivo* tumorigenic potential of BMs surgical samples correlates with patient poor survival. Data is expressed as median with interquartile range. Differences were considered statistically significant for p-values<0.05, according to the Kruskal-Wallis with Dunn’s multiple comparisons and Log-rank (Mantel-Cox) tests.

**Table 1.**
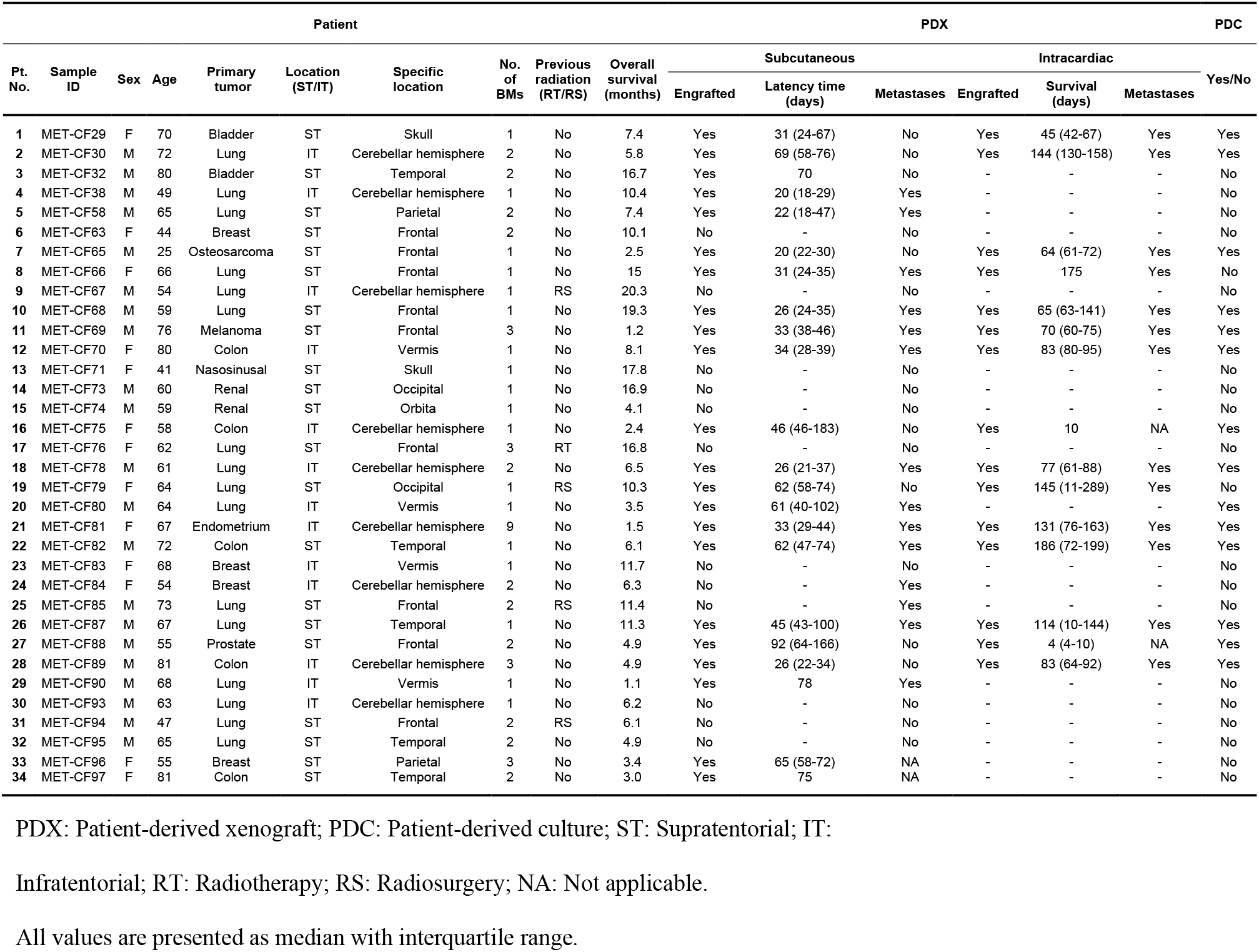
Summary of the clinical data of BMs patients and experimental results from PDX and PDC models.

The cohort of patients that generated the BMs tumor models had a median age of 64 years (25-81 years) (**Figure S1A**), with a male predominance (n=21; 62%) (**Figure S1B**). The most common primary tumor origins were lung (47%), colon (15%) and breast (12%) (**Figure S1C**), reflecting the tumor types with highest predisposition to originate BMs. Within the 16 lung cancer samples, 14 were classified as non-small cell carcinomas. Most BMs were located in the supratentorial compartment, in the frontal or temporal lobes (**Figures S1D–S1E**). Although all primary cancer types disseminated to the supratentorial compartment, only few were found in the posterior fossa (**Figures S1F-S1G**). The patients’ median survival after the diagnosis of BMs was 6.4 months (**Figure S1H**) and a trend towards a worse outcome was seen in patients with BMs within the infratentorial compartment (**Figure S1I**).

### *In vivo* tumorigenicity and clinical aggressiveness of subcutaneously implanted human brain metastases

Among multiple BMs samples, from 10 different primary cancer origins, implanted in the flank of NSG mice, we successfully grew tumors from 8 distinct cancer types. *In vivo* tumorigenicity was defined as tumor formation within 6 months after flank implantation, confirmed by histopathological analysis. The overall take rate for establishing PDXs of BMs was 65% (**Figure 1B**). Latency time, defined as the time between the subcutaneous implantation of the tumor fragment and the detection of a measurable lesion, decreased through passaging (**Figure 1C**). Notably, the ability of transplants to form tumors *in vivo* correlated with the donor patient’s clinical outcome. Engrafted BMs (n=22) derived from patients with an overall survival of 6 months, in contrast with BMs unable to originate tumors (n=12) whose patients had an overall survival of 10.75 months (**Figure 1D**; p=0.0021). A correlation between the *in vivo* tumorigenic potential and other clinical factors such as age, gender and location of the BMs was not observed (data not shown).

### Spontaneous dissemination of cancer cells in subcutaneous xenografts

The value of subcutaneous xenografts of BMs as a model to study human metastatic disease would be greatly increased if they recapitulated the dissemination pattern of the patient’s original tumor. To elucidate the metastatic potential of the implanted human BMs, we performed histopathological analysis of all the organs in each mouse. Remarkably, of the 22 tumors engrafted in the flank of NSG mice, 64% (n=14) originated spontaneous metastases to different sites, suggesting that cancer cells derived from BMs maintain their original metastatic potential. Moreover, we observed homing of cancer cells to the primary tumor site in 78% (7/9) of lung cancer BMs (**Figure 2A and Figure S2**). The sites of metastases across serial *in vivo* passaging varied with different human samples and occurred both at early and late passages. Interestingly, the spontaneous metastatic phenotype to the CNS (3/14; 21%) was observed in early passages and presented as leptomeningeal dissemination. The number of metastatic sites decreased with *in vivo* passaging (**Figure 2B**), possibly due to loss of heterogeneity by selection of more proliferative clones.

**Figure 2.**
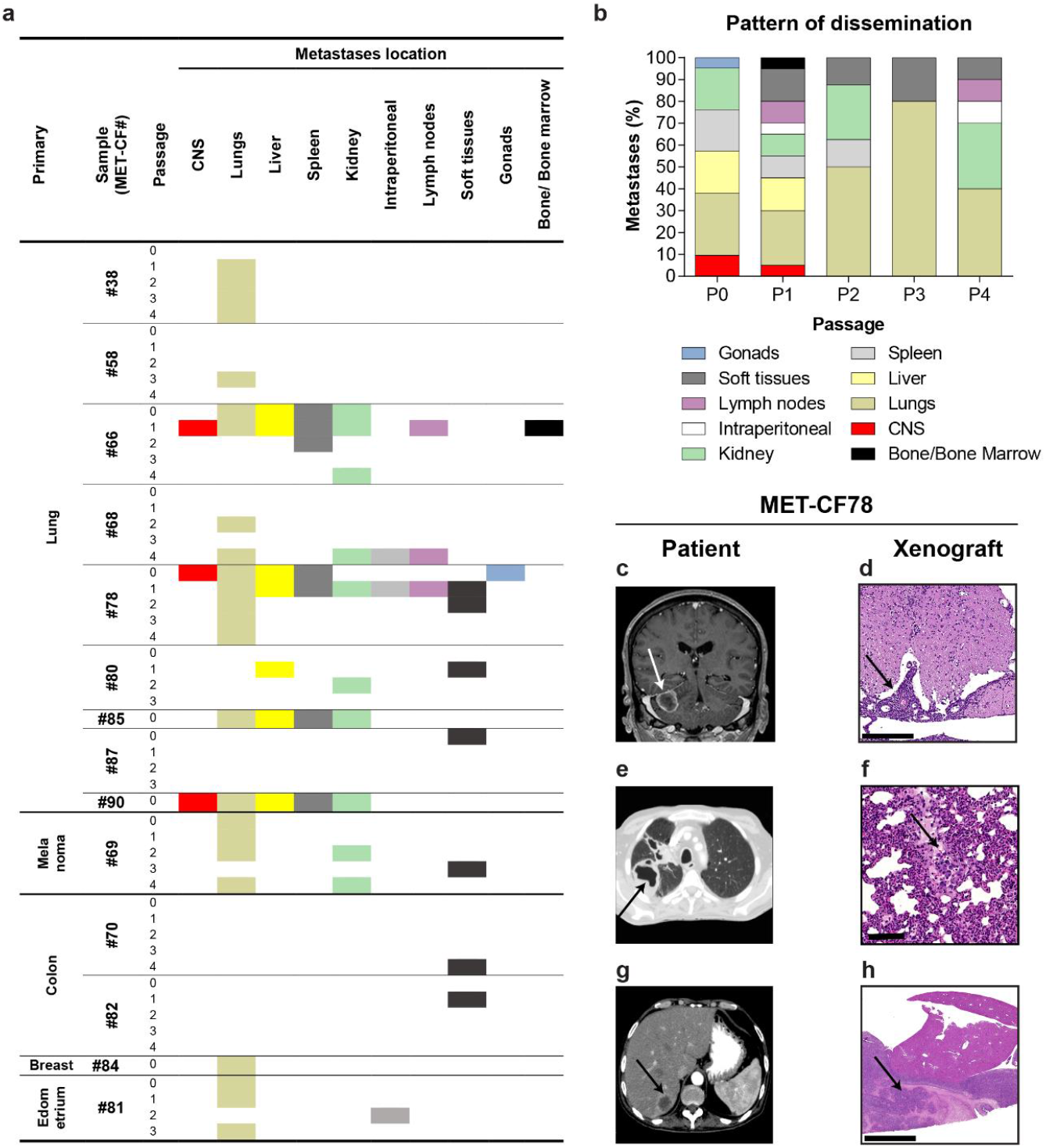
Spontaneous dissemination of subcutaneously implanted tumors. (**A**) Metastatic sites observed during human BMs expansion in mice. (**B**) Pattern of dissemination of cancer cells across *in vivo* passages. (**C-H**) Representative clinical case of a 61 years-old male patient with metastatic lung carcinoma to the brain and to the liver, whose PDX mimicked the donor patient disease. (**C**) Magnetic Resonance Imaging (MRI) of the brain, coronal T1 contrast-enhanced, showing a right cerebellar hemisphere BM. (**D**) H&E staining of the mouse lumbar spinal cord with leptomeningeal dissemination (arrow). (**E**) Computed Tomography (CT) scan of the patient’s thorax showing a primary lung cancer on the right lung and (**F**) H&E staining of the matched mouse lung with a metastasis (arrow). (**G**) Patient abdominal CT scan showing a liver metastasis (arrow) and (**H**) the matched liver metastasis in the xenograft.

We have compared the pattern of dissemination of each PDX with the patient whom the sample derived from. We found that the pattern of spontaneous cancer dissemination in some subcutaneous PDXs recapitulated the metastatic phenotype of the patients’ disease (**Table S1**). Mouse xenografts and matched patients shared metastatic sites in 7 samples (50%). For example, MET-CF78 exhibited metastatic deposits in the same organs as the patient with stage IV lung cancer from where it originated (**Figure 2C**). MET-CF69 was derived from a patient with metastatic melanoma and a diffuse lung infiltration, a pattern also observed in the corresponding xenograft (**Figure S3**). Importantly, three animal models (21%) showed spontaneous leptomeningeal dissemination to the CNS (**Figures 2C-2D and Figure S4**).

### Intracardiac PDXs increase the metastatic potential of cancer cells to CNS and mimic human disease

The injection of cancer cells in the left cardiac ventricle of mice has been previously validated as a good animal model for the study of BMs. We have isolated tumor cells from 15 BMs, which were consecutively passaged (up to passage 3) by intracardiac injection into NSG mice (**Figure 3A**). The most common primary tumors were lung cancer (40%) and colon cancer (27%). Mouse survival decreased with passaging (**Figure 3A, B**), likely reflecting the selection of more aggressive cancer cells. The survival of the animals also varied according to the primary tumor type (**Figure 3C**) and the location of the BMs in the intracranial compartment (**Figure S5**). Interestingly, intracardiac PDXs derived from infratentorial metastases showed a trend towards a worse outcome, mimicking the BMs patient cohort (**Figure S1I**).

**Figure 3.**
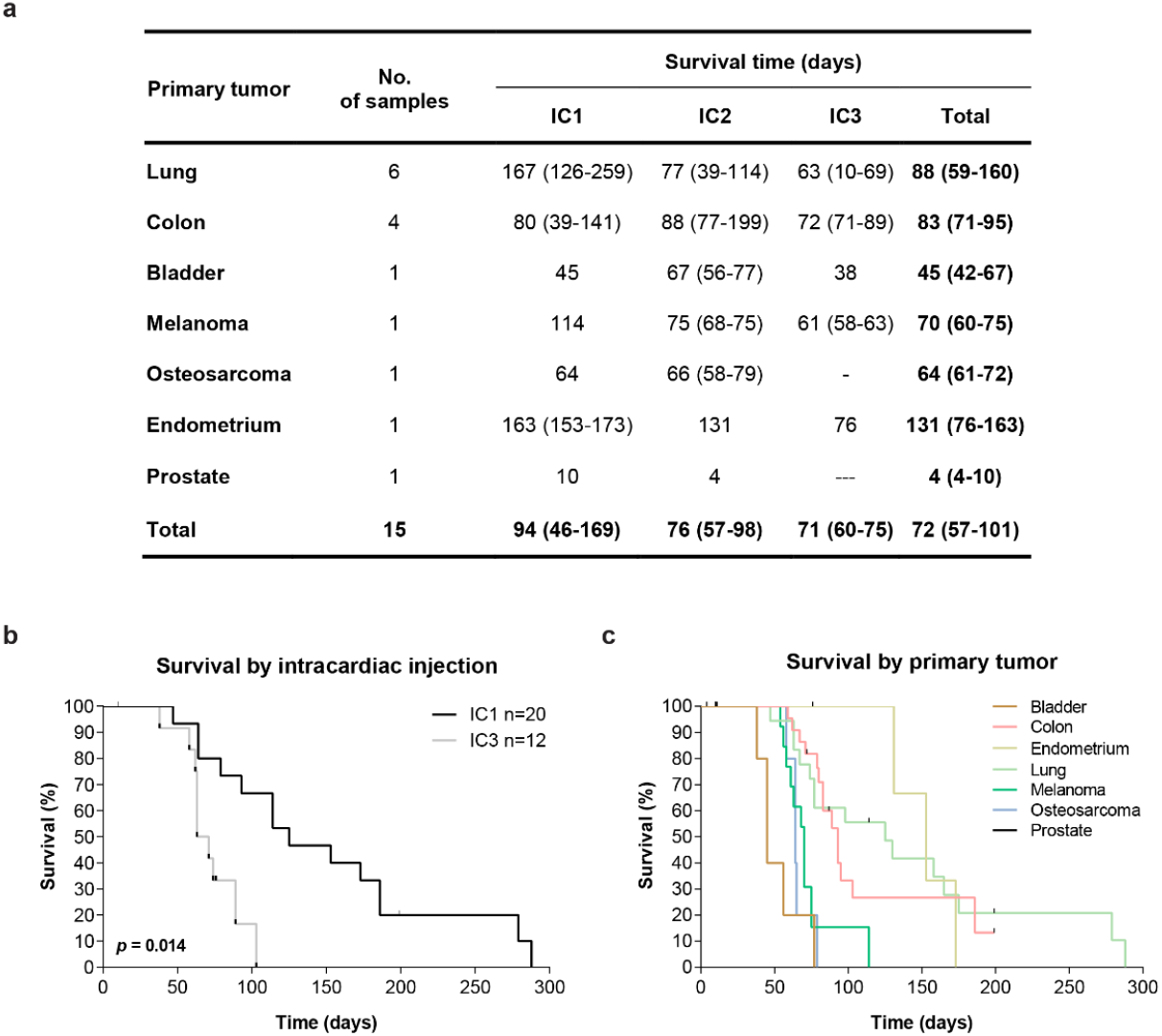
Intracardiac xenografts derived from human BMs samples. (**A**) Survival of mice submitted to intracardiac injections of cancer cells derived from human BMs of diverse primary cancers. (**B**) Kaplan-Meier survival curves in early (IC1) and late (IC3) intracardiac xenografts. (**C**) Mice overall survival according to the primary tumor. Data is expressed as median with interquartile range. Differences were considered statistically significant for p-values<0.05, according to the Log-rank (Mantel-Cox) test.

All the intracardiac PDXs we analyzed developed systemic metastases. As expected from the delivery of cancer cells into the blood stream, the number of metastatic sites increased when compared to subcutaneous tumor implantation (**Figure 4A**). The intracardiac model also increased the tropism of cancer cells to the CNS (6/13; 46%), either focal metastases in the parenchyma or leptomeningeal dissemination, particularly at later passages (**Figure 4B**).

**Figure 4.**
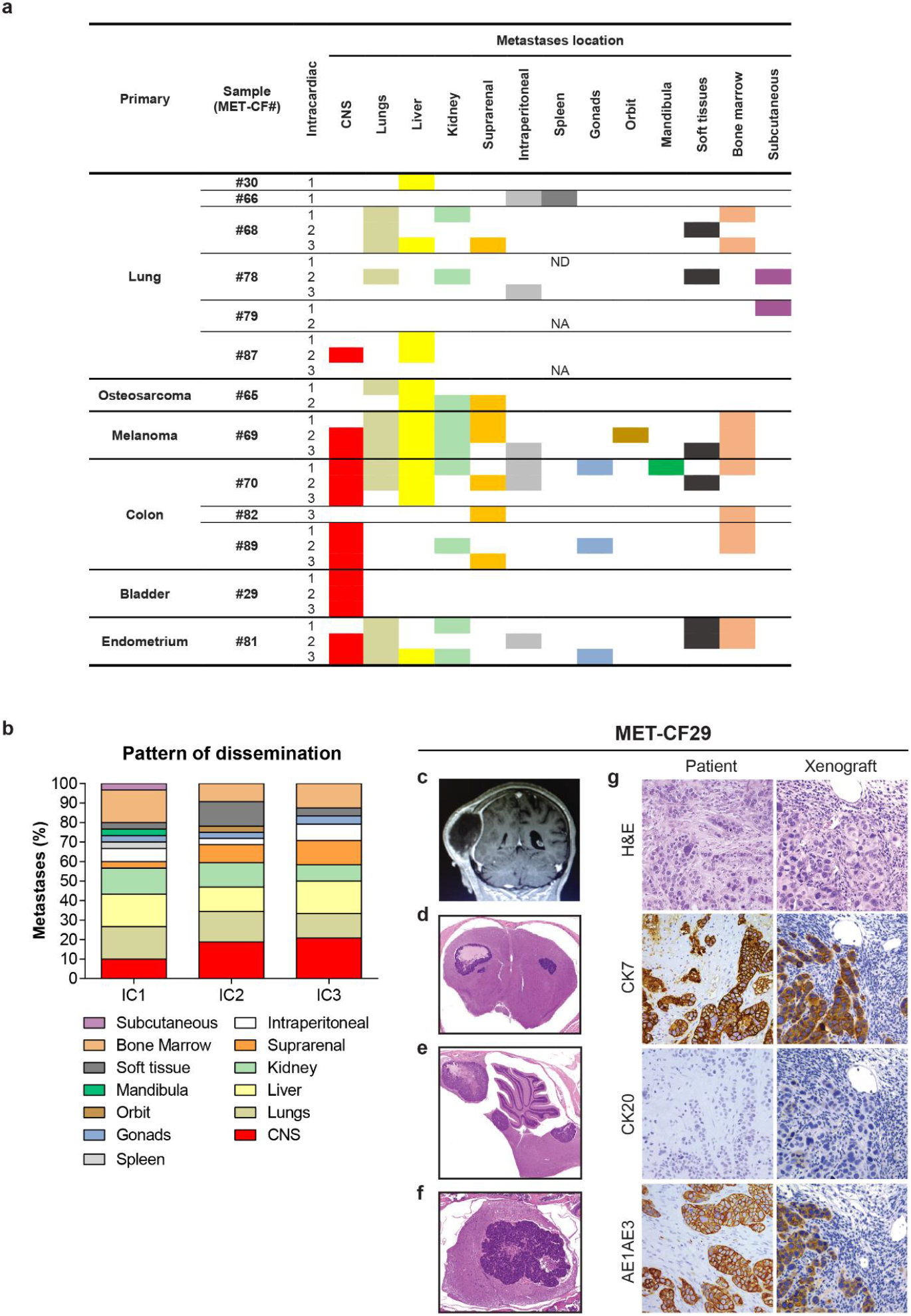
Metastatic phenotype in intracardiac xenograft models of human BMs. (**A**) Systemic location of metastases during serial injection of human BMs in mice. (**B**) Pattern of dissemination of cancer cells upon intracardiac injections. (**C-G**) Intracardiac mouse xenograft from a 70 years-old female patient with a bladder carcinoma. Contrast-enhanced coronal T1 MRI sequence of a parietal skull metastasis with adjacent dural and subcutaneous tissue invasion (**C**). Representative H&E stained sections of the correspondent mouse xenograft revealing exclusive CNS metastases, in the supratentorial compartment (**D**), in the infratentorial compartment (**E**) and in the spinal cord (**F**). Comparison of the immunohistochemical markers for bladder carcinoma between human parental BM sample and corresponding xenografted tumor (**G**).

When we compared the pattern of cancer cell dissemination in the intracardiac xenograft models with the respective patient metastatic disease (**Table S2**), we observed that 8 PDXs (62%) disseminated to the same organs as their respective donor patients and, interestingly 3 PDXs (23%) presented metastases in the organ correspondent to the primary tumor. (**Figure S6 and S7**). Of notice, MET-CF29, derived from a patient with metastatic bladder carcinoma, exhibited metastases exclusive to the CNS and the same pattern of immunostaining as the corresponding patients’ BM (**Figures 4C-G**).

Altogether, our results demonstrate that intracardiac PDXs of BMs recapitulate human metastatic disease and may constitute good models to study CNS dissemination from diverse primary cancers.

## Effective therapeutic response of patient-derived models to targeted anticancer drugs

To assess the potential of our patient-derived models in evaluating the response to targeted therapies, we preclinically tested the FDA approved drugs buparlisib (pan-PI3K inhibitor) and everolimus (mTOR inhibitor), targeting the PI3K/AKT/mTOR pathway. These drugs have previously demonstrated efficacy in orthotopic models of BMs from breast cancer[55]. Moreover, it has been reported that BMs have genomic alterations associated with sensitivity to PI3K/AKT/mTOR inhibitors[21]. Both compounds effectively inhibited their expected targets at low concentrations in a PDC of a lung cancer BM (MET-CF78). This is shown by reduced AKT and S6 phosphorylation upon buparlisib treatment and by downregulation of S6, but not AKT phosphorylation in the case of everolimus (**Figure 5A**). Accordingly, both drugs significantly reduced cell proliferation in a dose-dependent manner (**Figure 5B-5C**).

**Figure 5.**
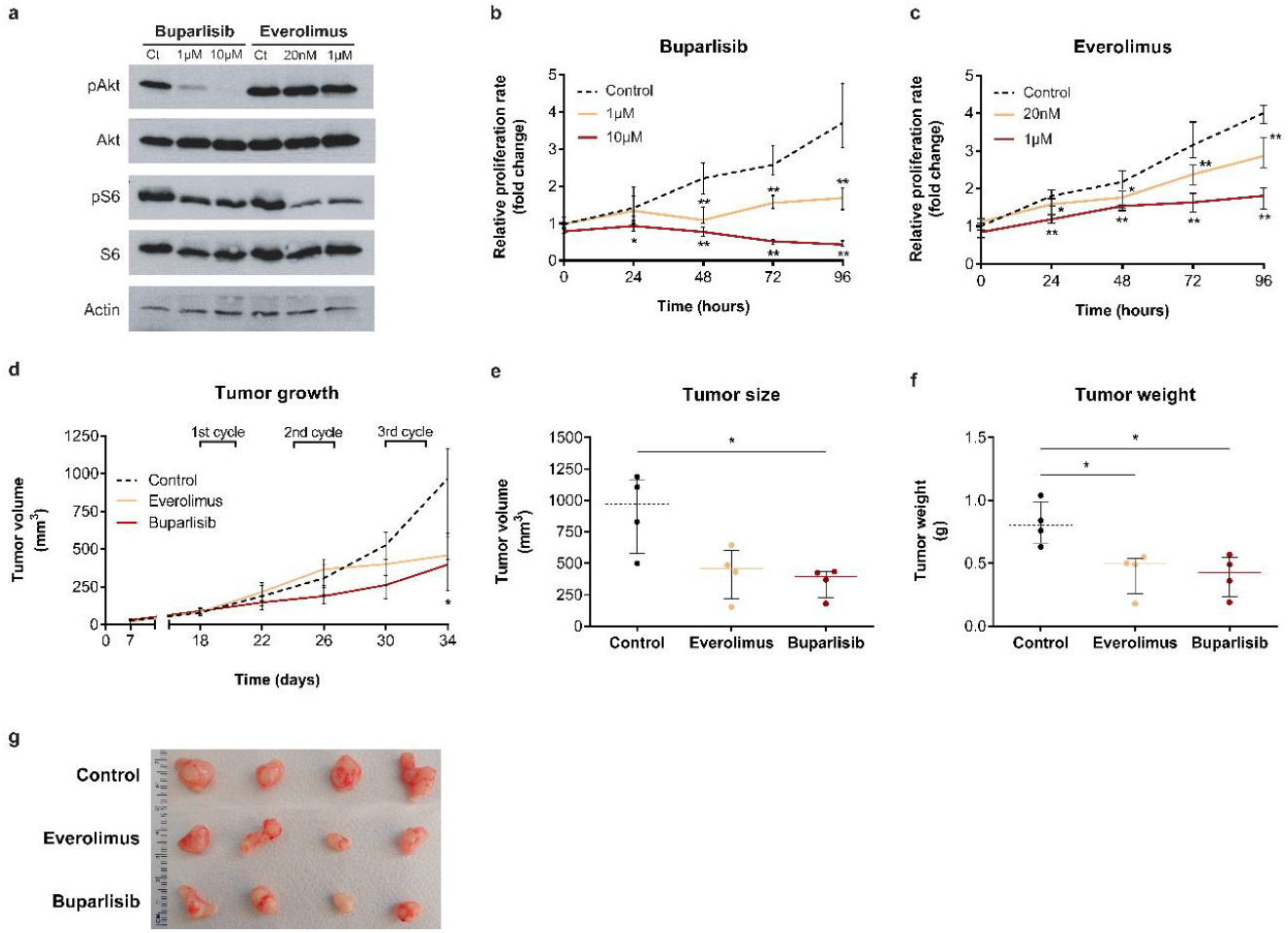
Patient-derived models of lung cancer BMs respond to PI3K and mTOR inhibition. (**A**) Representative Western blot demonstrating AKT and mTOR pathway inhibition by buparlisib and everolimus, respectively. Inhibition of cell proliferation upon treatment with increasing doses of buparlisib (**B**) and everolimus (**C**). Cell viability was measured by MTS assay. Data represents median of triplicates with interquartile range. (**D**) Inhibition of flank tumor growth upon three cycles of therapy with buparlisib and everolimus. Significant reduction in tumor size (**E**) and tumor weight (**F**) by the end of treatment. (**G**) Representative photographs of flank tumors in each experimental group by the end of treatment.

To further evaluate the utility of our models for precision medicine, we treated the subcutaneous PDX of MET-CF78 with three cycles of oral therapy with buparlisib or everolimus (four days on and 2 days off). We have used a subcutaneous model to better assess treatment efficacy in vivo. Both compounds effectively reduced tumor growth (**Figure 5D**) and, by the end of the therapeutic protocol, tumor size and tumor weight were significantly lower in the treated tumors (**Figure 5E-5G**). Our patient-derived models provide a relevant tool for preclinical testing of anticancer therapies.

## DISCUSSION

PDX models have recently emerged has good preclinical platforms for drug discovery in oncology[32, 57]. However, the development of PDXs from BMs has been limited to few studies using samples from the most common cancer types (breast and lung cancers). Contreras-Zarate *et al.* successfully generated 8 PDXs of breast cancer BMs upon implantation in the mammary fat pad[54]. Lee *et al.* established subcutaneous and orthotopic PDXs from non-small cell lung cancer (NSCLC) primary tumors and BMs, which maintained the histopathological similarities and the molecular profiling signatures of the parental tumors. The intracardiac injection of three primary cultured human cancer cells in mice originated systemic tumors but failed to recapitulate the metastatic phenotype of the patients’ disease[41]. Using intracranial implantation of patient-derived BMs tumorspheres from lung cancer, Nolte *el al.* showed that those cells could reproduce the original patients’ tumors[58]. A recent study described spontaneous metastases in melanoma subcutaneous PDXs but no correspondence with patients’ disease was reported[59]. To our knowledge, our study is the first to describe the full spectrum of human’s metastatic disease mirrored in subcutaneous and intracardiac xenograft models of BMs. Furthermore, we demonstrate the value of these models for preclinical testing of targeted anticancer therapies.

We have built a unique collection of 22 well-established PDXs of BMs from 8 distinct primary cancers, which were phenotypically characterized over time and compared to the original patients’ metastatic disease. A noteworthy observation from our subcutaneous xenograft models is their capability to spontaneously disseminate to distinct organs in mice, namely to the location of the patients’ primary tumor and to the CNS where the surgical sample was collected. More importantly, mouse subcutaneous xenografts and their human counterpart shared metastatic sites in 7 samples (50%) reinforcing the relevance of our model to study human disease.

The intracardiac injection of cancer cells has been extensively studied and validated as a good model to study BMs. However, the published studies relied on immortalized cancer cells derived from murine[23] or human[20] cell lines, generated through successive rounds of intracardiac injection in mice for the selection of brain-tropic cells. The rapid progression of the brain metastatic disease in these models limits their utility for drug testing and, more importantly, they do not fully portray the heterogeneity of human tumors. The systematic intracardiac injection of patient-derived brain metastatic cells and the thorough assessment of cancer cell fate in different organs have not been addressed. The results from our study have identified advantages of the intracardiac PDX model for the study of human metastatic cancers when compared to the subcutaneous model including: a) high incidence of spontaneous metastases (100% in intracardiac PDXs vs 64% in subcutaneous PDXs); b) increased CNS dissemination (46% in intracardiac PDXs vs 21% in subcutaneous PDXs); and c) higher number of metastatic sites shared between xenografts and donor patients (62% in intracardiac PDXs vs 50% in subcutaneous PDXs). Importantly, the library of xenograft-matched PDCs we generated constitutes an excellent tool for streamlined drug discovery: *in vitro* high-throughput drug screens to identify promising compounds can readily advance into *in vivo* preclinical trials. Together, these features make our patient-derived models of BMs suitable to study the biology of human metastatic cancers and to perform preclinical drug testing of patient-tailored therapies, which we illustrated with the use of buparlisib and everolimus.

Recent studies using mouse models of cancer provided evidence of polyclonal seeding of cancer cells, which often disseminate in parallel to form metastases, and of cooperation between subclones to enhance tumor progression[60-63]. Phylogenetic analysis of matched samples from metastatic cancer patients corroborated those findings and demonstrated that metastasis-to-metastasis spread is a common event[64]. These mechanisms likely enhance tumor heterogeneity and may potentially contribute to drug resistance. Although future genomic studies need to confirm the clonal identity of each metastatic site in our PDXs, the observed pattern of cancer dissemination over serial passaging in mice, particularly with the intracardiac injection, might reflect the conservation of inter- and intra-tumoral heterogeneity. To develop our xenografts, we either used tumor fragments or dissociated cancer cells from fragments of subcutaneously grown BMs, thereby avoiding artificial clonal selection.

The absence of mouse models that accurately reflect the human metastatic disease to the brain has been a barrier to the investigation of disease drivers and to the identification of site-specific targeted treatments. Since cancer patients’ survival is dramatically decreased upon the establishment of BMs, the development of such tools is an urgent need. Our translational models (using BMs of diverse primary cancer origins that recapitulate the dynamics of cancer cell dissemination, mirror patient metastatic disease and respond to anticancer therapies) represent outstanding tools to advance the understanding of the biology of BMs and to foster the rapid discovery of novel therapeutics.

## MATERIALS AND METHODS

### Specimen collection, annotation and biobanking

Samples were collected in accordance with the Hospital de Santa Maria (Centro Hospitalar Universitário Lisboa Norte, CHULN, Lisbon, Portugal) Ethics Board and a written informed consent was obtained from all patients prior to study participation. Specimens and detailed clinical annotations were obtained from 34 patients with BMs from diverse primary tumors submitted to surgical resection at the Department of Neurosurgery of Hospital de Santa Maria (CHLN, Lisbon, Portugal) from 2015 to 2017. Surgical BMs samples not needed for diagnostic purposes were divided into four portions within 1 hour after surgery: for implantation into immune-deficient mice, DNA/RNA extraction, pathologic assessment, and storage. Samples were stored at Biobanco-iMM CAML (Biobank of the Lisbon Academic Medical Center, Lisbon, Portugal).

### Establishment of subcutaneous patient-derived xenografts (PDXs) of BMs

In accordance with Directive 2010/63/EU (transposed to Portuguese legislation through Decreto-Lei No. 113/2013, of August 7th), all animal procedures were approved by the institutional animal welfare body (ORBEA-iMM), in order to ensure that the use of animals complies with all applicable legislation and following the 3R’s principle, as well as licensed by the Portuguese competent authority (license number: 012028\2016). All animals were kept in specific pathogen-free (SPF) conditions, randomly housed per groups under standard laboratory conditions (at 20-22°C under 10 hour light/14 hour dark), and given free access to food (RM3, SDS Diets, Witham, UK) and water (Ultrapure). Invasive procedures were performed with animals under volatile anesthesia (isoflurane) administered using a RC^2^ Rodent Circuit Controller Anesthesia System (VetEquip®, California, USA). Humane endpoints were established for tumor volume of 1000mm^3^, 10% body weight loss, paralysis and abdominal distension. Animals were euthanized using anesthetic overdose.

NOD.Cg-*Prkdc^scid^ Il2rg^tm1Wjl^/SzJ* (NSG) mice were purchased from Charles River Laboratories (Massachusetts, USA) and used as hosts for PDXs production. Human BMs samples were divided into small fragments (ca. 4×4×4mm) and implanted into the subcutaneous tissue of each mouse flank (1 fragment per mouse, n=2 mice/sample/passage). All mice were monitored for tumor growth, body weight, discomfort and distress every other day. Once palpable, tumor size was measured twice a week using a caliper (VWR, Pennsylvania, USA), and the volume was estimated using the formula (AxB^2^)/2 where A is the antero-posterior measure and B is the dorsoventral measure. When tumors reached 1000mm^3^, mice were euthanized, the flank tumors were removed and serially passaged *in vivo.* Histopathologic analysis was also performed in other organs, namely CNS, femur and tibia, liver, lungs, spleen, pancreas, adrenal and reproductive systems, lymph nodes, thymus, heart and kidneys. Tumors were serially passaged until passage (P) 4.

### Establishment of patient-derived cultures (PDCs)

Freshly harvested PDX tumors were minced using sterile scalpel blades and enzymatically dissociated using trypsin, hyaluronidase, kynurenic acid, DNase (Sigma-Aldrich, Missouri, USA) in artificial cerebrospinal fluid, at 37°C with agitation for ~30min. Enzymatic dissociation was halted by adding trypsin inhibitor (Sigma-Aldrich, Missouri, USA). Cell suspension was further centrifuged at 1000rpm for 5min. Supernatant was then discarded, pelleted cells were washed in DMEM/F12 (Gibco, California, USA) and filtered through a 70μm cell strainer. Dissociated cells were cultured in DMEM/F12 supplemented with 10% FBS (Biochrom, Berlin, Germany), 2% B27, 1% HEPES, 1% L-Glutamine and 1x Antibiotic-Antimycotic (Gibco, California, USA) at 37°C with 5% CO_2_. Growth factors were supplemented every 2/3 days.

### Establishment of intracardiac PDXs of BMs

Freshly dissociated viable cells were counted using trypan blue exclusion and 50,000 cells re-suspended in 80μl PBS were injected in the left cardiac ventricle of recipient NSG mice (n=2/sample/passage). Intracardiac injections were performed under volatile anesthesia (Isoflurane). After injection mice recovered on a heating pad while being observed for adverse events, such as cardio-respiratory failure. All mice were monitored for neurological disabilities, body weight, discomfort and distress every other day. Mice were euthanized when presented more than one of these manifestations and the organs were collected for histopathological analysis, as described above.

### Histological and immunohistochemical analysis

Tissue samples were fixed immediately in 10% neutral buffered formalin solution, dehydrated and embedded in paraffin, serially sectioned at a thickness of 4μm using a microtome, mounted on microscope slides and stained with hematoxylin and eosin (H&E) for morphological examination. Tissue sections were also used for immunohistochemical (IHC) staining following conventional protocols, using the following primary antibodies: CK7 (Invitrogen 180234, California, USA, Thermo Fisher Scientific Cat# MA1-80234, RRID:AB_928715), CK20 (Dako Hs20.8, California, USA, Agilent Cat# M7019, RRID:AB_2133718), AE1AE3 (Invitrogen 180132, California, USA, Thermo Fisher Scientific Cat# MA5-13203, RRID:AB_10942225), and mitochondria (Merck MAB1273, Massachusetts, USA, Millipore Cat# MAB1273, RRID:AB_9405). Briefly, for antigen retrieval slides were treated in a PT Link module (DAKO, California, USA) at low-Ph, followed by incubation with the primary antibodies. EnVision Link horseradish peroxidase/DAB visualization system (Dako, California, USA) was used, and sections were then counterstained with hematoxylin and mounted. H&E and IHC slides were examined by a specialized pathologist, and representative photomicrographs were taken using Leica DM2500 microscope coupled to a Leica MC170 HD microscope camera (Wetzlar, Germany).

### *In vitro* drug assay

PDCs from MET-CF78 BM sample, were seeded in 96-well plates, 1000 cells per well, and incubated up to 96h with either 1μM or 10μM buparlisib (NVP-BKM120 PI3K inhibitor, Selleckchem, Munich, Germany), and with either 20nM or 1μM of everolimus (RAD001, mTOR inhibitor, Selleckchem, Munich, Germany).

Cell viability was determined by MTS (CellTiter 96 Aqueous One Solution Reagent, Promega, Wisconsin, USA) as previously described[65]. Briefly, cells were incubated for 2h with MTS and the absorbance was measured at 490nm. Two independent experiments were performed with 3 technical repetitions each.

For Western blot analysis, cells were seeded in 6-well plates at 0.5 x 10^6^ cells per well and incubated for 2h with the above-mentioned drug conditions. Cell lysates used for immunoblotting were prepared as previously described[66]. Cells were lysed in lysis buffer supplemented with 1x phosphatase inhibitors (PhosStop, Roche Diagnostics, Basel, Switzerland) and a 1x protease inhibitor cocktail (Complete Mini, Roche Diagnostics, Basel, Switzerland). After centrifugation at 10,000g for 15 minutes at 4°C, the supernatant was harvested. Protein concentration was determined using the Bradford protein assay (BioRad, California, USA). Equal amounts of protein were subjected to SDS-PAGE and transferred onto nitrocellulose membranes (BioRad, California, USA), which were blocked with 5% skim milk for 1 hour at room temperature, incubated with the primary antibodies overnight, and then blotted with the appropriate secondary antibodies. Antibodies against p-S473-AKT (Cell Signaling Technology Cat# 4060, RRID:AB_231504), AKT (Cell Signaling Technology Cat# 9272, RRID:AB_329827), p-S235/236-S6 (Cell Signaling Technology Cat# 2211, RRID:AB_331679) and S6 (Cell Signaling Technology Cat# 2217, RRID:AB_331355, Massachusetts, USA) and actin (Santa Cruz Biotechnology Cat# sc-47778 HRP, RRID:AB_2714189, California, USA,) were used[67]. Two independent experiments were performed.

### *In vivo* drug treatment of subcutaneous PDX

*In vivo* drug response evaluation was performed using NSG mice injected subcutaneously in the flank with MET-CF78 cells (1.5×10^6^ cells/mouse). The MET-CF78 culture was derived from a patient with lung cancer BM. When all tumors were measurable (50-150 mm^3^), animals were randomized into three treatment groups (N=7 mice/group): untreated control (DMSO/PEG400), mTOR inhibitor everolimus (3mg/kg/day) and PI3K inhibitor buparlisib (30mg/kg/day). Both compounds were purchased from Selleck Chemicals (Munich, Germany), dissolved in dimethyl sulfoxide (DMSO, Sigma-Aldrich, Darmstadt, Germany) and stored at −80°C. Drugs were freshly diluted in polyethylene glycol 400 (PEG400, Sigma-Aldrich, Darmstadt, Germany) (30% v/v) immediately before administration by oral gavage. Mice received three cycles of therapy (four days on and two days off). The animals were monitored daily, body weight variations were recorded throughout treatment, and tumor volume was measured using a caliper. Mice were euthanized by the end of the treatment (day 34) and tumor samples were collected for histopathological analysis, as described above.

### Statistical analysis

Statistical differences were determined with non-parametric Kruskal-Wallis (Dunn’s Multiple Comparison tests) and Mann–Whitney tests using GraphPad Prism v6.0 (GraphPad, California, USA, GraphPad Prism, RRID:SCR_002798). Survival curves were analyzed using log-rank tests (Mantel-Cox). Differences were considered statistically significant for p<0.05.

## ACKNOWLEDGMENTS

The authors would like to acknowledge the patients who kindly provided the tumor specimens used to generate the PDXs models needed for this research. The Biobanco-iMM CAML enabled the process of tumor specimen collection, processing and storage. Finally, authors would like to acknowledge the Histology and Comparative Pathology Laboratory and the Rodent Facility from Instituto de Medicina Molecular João Lobo Antunes for technical assistance.

## AUTHOR CONTRIBUTIONS

Study design: CF, RC and JTB. Study conduct: CC, RC, EP, TC, PP and RR. Data collection: CF, CC, EP, TC, PP and RR. Data analysis: CF, CC, RC, EP and TC. Data interpretation: CF, CC, RC, EP, TC and JTB. Drafting manuscript: CF and RC. Revising manuscript content: CF, CC, RC, EP, TC, PP, RR, JP, JM and JTB. Approving final version of manuscript: CF, CC, RC, EP, TC, PP, RR, JP, JM and JTB. CF, CC, RC, EP, TC, PP, RR, JP, JM and JTB take responsibility for the integrity of the data analysis.

## DATA AND MATERIALS AVAILABILITY

All data associated with this study are available in the main text or the supplementary materials.

## FUNDING

CC was supported by a fellowship from Fundação para a Ciência e a Tecnologia (FCT, SFRH/BD/140299/2018). EP was supported by a fellowship from FCT (PD/BD/128288/2017). This project was funded by FCT (PTDC/MED-ONC/32222/2017), Fundação Millennium bcp and by private donations. This work was also supported by UID/BIM/50005/2019, co-funded by FCT/Ministério da Ciência, Tecnologia e Ensino Superior (MCTES) through funds of Programa de Investimento e Despesas de Desenvolvimento da Administração Central (PIDDAC). The funders had no role in study design, data collection and analysis, decision to publish, or preparation of the manuscript.

## CONFLICTS OF INTEREST

The authors have declared that no competing interests exist.

## Notes

### Competing Interest Statement

The authors have declared no competing interest.

## REFERENCES

1. Gavrilovic, I.T. and J.B. Posner, Brain metastases: epidemiology and pathophysiology. J Neurooncol, 2005. 75(1): p. 5–14.

2. Achrol, A.S., et al., Brain metastases. Nat Rev Dis Primers, 2019. 5(1): p. 5.

3. Glantz, M.J., et al., Diagnosis, management, and survival of patients with leptomeningeal cancer based on cerebrospinal fluid-flow status. Cancer, 1995. 75(12): p. 2919–31.

4. Taillibert, S. and M.C. Chamberlain, Leptomeningeal metastasis. Handb Clin Neurol, 2018. 149: p. 169–204.

5. Leal, T., et al., Leptomeningeal Metastasis: Challenges in Diagnosis and Treatment. Curr Cancer Ther Rev, 2011. 7(4): p. 319–327.

6. Soffietti, R., R. Ruda, and R. Mutani, Management of brain metastases. J Neurol, 2002. 249(10): p. 1357–69.

7. Bhambhvani, H.P., et al., The primary sites leading to brain metastases: Shifting trends at a tertiary care center. J Clin Neurosci, 2020. 80: p. 121–124.

8. Nayak, L., E.Q. Lee, and P.Y. Wen, Epidemiology of brain metastases. Curr Oncol Rep, 2012. 14(1): p. 48–54.

9. Barnholtz-Sloan, J.S., et al., Incidence proportions of brain metastases in patients diagnosed (1973 to 2001) in the Metropolitan Detroit Cancer Surveillance System. J Clin Oncol, 2004. 22(14): p. 2865–72.

10. Sperduto, P.W., et al., Diagnosis-specific prognostic factors, indexes, and treatment outcomes for patients with newly diagnosed brain metastases: a multi-institutional analysis of 4,259 patients. Int J Radiat Oncol Biol Phys, 2010. 77(3): p. 655–61.

11. Berghoff, A.S., et al., Descriptive statistical analysis of a real life cohort of 2419 patients with brain metastases of solid cancers. ESMO Open, 2016. 1(2): p. e000024.

12. Lin, N.U., J.R. Bellon, and E.P. Winer, CNS metastases in breast cancer. J Clin Oncol, 2004. 22(17): p. 3608–17.

13. Lockman, P.R., et al., Heterogeneous blood-tumor barrier permeability determines drug efficacy in experimental brain metastases of breast cancer. Clin Cancer Res, 2010. 16(23): p. 5664–78.

14. Groves, M.D., New strategies in the management of leptomeningeal metastases. Arch Neurol, 2010. 67(3): p. 305–12.

15. Langley, R.R. and I.J. Fidler, The biology of brain metastasis. Clin Chem, 2013. 59(1): p. 180–9.

16. Lin, X. and L.M. DeAngelis, Treatment of Brain Metastases. J Clin Oncol, 2015. 33(30): p. 3475–84.

17. Pitz, M.W., et al., Tissue concentration of systemically administered antineoplastic agents in human brain tumors. J Neurooncol, 2011. 104(3): p. 629–38.

18. Heerboth, S., et al., EMT and tumor metastasis. Clin Transl Med, 2015. 4: p. 6.

19. Venur, V.A., U.N. Chukwueke, and E.Q. Lee, Advances in Management of Brain and Leptomeningeal Metastases. Curr Neurol Neurosci Rep, 2020. 20(7): p. 26.

20. Bos, P.D., et al., Genes that mediate breast cancer metastasis to the brain. Nature, 2009. 459(7249): p. 1005–9.

21. Brastianos, P.K., et al., Genomic Characterization of Brain Metastases Reveals Branched Evolution and Potential Therapeutic Targets. Cancer Discov, 2015. 5(11): p. 1164–1177.

22. Richichi, C., et al., Mutations targeting the coagulation pathway are enriched in brain metastases. Sci Rep, 2017. 7(1): p. 6573.

23. Valiente, M., et al., Serpins promote cancer cell survival and vascular co-option in brain metastasis. Cell, 2014. 156(5): p. 1002–16.

24. Rabbie, R., et al., The mutational landscape of melanoma brain metastases presenting as the first visceral site of recurrence. Br J Cancer, 2020.

25. Tentler, J.J., et al., Patient-derived tumour xenografts as models for oncology drug development. Nat Rev Clin Oncol, 2012. 9(6): p. 338–50.

26. Rosfjord, E., et al., Advances in patient-derived tumor xenografts: from target identification to predicting clinical response rates in oncology. Biochem Pharmacol, 2014. 91(2): p. 135–43.

27. Hidalgo, M., et al., Patient-derived xenograft models: an emerging platform for translational cancer research. Cancer Discov, 2014. 4(9): p. 998–1013.

28. To, B., D. Isaac, and E.R. Andrechek, Studying Lymphatic Metastasis in Breast Cancer: Current Models, Strategies, and Clinical Perspectives. J Mammary Gland Biol Neoplasia, 2020. 25(3): p. 191–203.

29. Murata, T., et al., Co-implantation of Tumor and Extensive Surrounding Tissue Improved the Establishment Rate of Surgical Specimens of Human-Patient Cancer in Nude Mice: Toward the Goal of Universal Individualized Cancer Therapy. In Vivo, 2020. 34(6): p. 3241–3245.

30. Sanden, E., et al., Establishment and characterization of an orthotopic patient-derived Group 3 medulloblastoma model for preclinical drug evaluation. Sci Rep, 2017. 7: p. 46366.

31. Rygaard, J. and C.O. Povlsen, Heterotransplantation of a human malignant tumour to “Nude” mice. Acta Pathol Microbiol Scand, 1969. 77(4): p. 758–60.

32. Bruna, A., et al., A Biobank of Breast Cancer Explants with Preserved Intra-tumor Heterogeneity to Screen Anticancer Compounds. Cell, 2016. 167(1): p. 260–274 e22.

33. Rubio-Viqueira, B., et al., An in vivo platform for translational drug development in pancreatic cancer. Clin Cancer Res, 2006. 12(15): p. 4652–61.

34. Siolas, D. and G.J. Hannon, Patient-derived tumor xenografts: transforming clinical samples into mouse models. Cancer Res, 2013. 73(17): p. 5315–9.

35. Braekeveldt, N., et al., Neuroblastoma patient-derived orthotopic xenografts retain metastatic patterns and geno- and phenotypes of patient tumours. Int J Cancer, 2015. 136(5): p. E252–61.

36. Joo, K.M., et al., Patient-specific orthotopic glioblastoma xenograft models recapitulate the histopathology and biology of human glioblastomas in situ. Cell Rep, 2013. 3(1): p. 260–73.

37. Choi, Y.Y., et al., Establishment and characterisation of patient-derived xenografts as paraclinical models for gastric cancer. Sci Rep, 2016. 6: p. 22172.

38. du Manoir, S., et al., Breast tumor PDXs are genetically plastic and correspond to a subset of aggressive cancers prone to relapse. Mol Oncol, 2014. 8(2): p. 431–43.

39. Cottu, P., et al., Modeling of response to endocrine therapy in a panel of human luminal breast cancer xenografts. Breast Cancer Res Treat, 2012. 133(2): p. 595–606.

40. Charafe-Jauffret, E., et al., ALDH1-positive cancer stem cells predict engraftment of primary breast tumors and are governed by a common stem cell program. Cancer Res, 2013. 73(24): p. 7290–300.

41. Lee, H.W., et al., Patient-derived xenografts from non-small cell lung cancer brain metastases are valuable translational platforms for the development of personalized targeted therapy. Clin Cancer Res, 2015. 21(5): p. 1172–82.

42. Dong, X., et al., Patient-derived first generation xenografts of non-small cell lung cancers: promising tools for predicting drug responses for personalized chemotherapy. Clin Cancer Res, 2010. 16(5): p. 1442–51.

43. John, T., et al., The ability to form primary tumor xenografts is predictive of increased risk of disease recurrence in early-stage non-small cell lung cancer. Clin Cancer Res, 2011. 17(1): p. 134–41.

44. Feldmann, G., et al., Cyclin-dependent kinase inhibitor Dinaciclib (SCH727965) inhibits pancreatic cancer growth and progression in murine xenograft models. Cancer Biol Ther, 2011. 12(7): p. 598–609.

45. Allaway, R.J., et al., Genomic characterization of patient-derived xenograft models established from fine needle aspirate biopsies of a primary pancreatic ductal adenocarcinoma and from patient-matched metastatic sites. Oncotarget, 2016. 7(13): p. 17087–102.

46. Villarroel, M.C., et al., Personalizing cancer treatment in the age of global genomic analyses: PALB2 gene mutations and the response to DNA damaging agents in pancreatic cancer. Mol Cancer Ther, 2011. 10(1): p. 3–8.

47. Fu, X.Y., et al., Models of human metastatic colon cancer in nude mice orthotopically constructed by using histologically intact patient specimens. Proc Natl Acad Sci U S A, 1991. 88(20): p. 9345–9.

48. Metildi, C.A., et al., Fluorescently labeled chimeric anti-CEA antibody improves detection and resection of human colon cancer in a patient-derived orthotopic xenograft (PDOX) nude mouse model. J Surg Oncol, 2014. 109(5): p. 451–8.

49. Delyon, J., et al., Validation of a preclinical model for assessment of drug efficacy in melanoma. Oncotarget, 2016. 7(11): p. 13069–81.

50. Colombo, P.E., et al., Ovarian carcinoma patient derived xenografts reproduce their tumor of origin and preserve an oligoclonal structure. Oncotarget, 2015. 6(29): p. 28327–40.

51. Ricci, F., et al., Patient-derived ovarian tumor xenografts recapitulate human clinicopathology and genetic alterations. Cancer Res, 2014. 74(23): p. 6980–90.

52. Karam, J.A., et al., Development and characterization of clinically relevant tumor models from patients with renal cell carcinoma. Eur Urol, 2011. 59(4): p. 619–28.

53. Sivanand, S., et al., A validated tumorgraft model reveals activity of dovitinib against renal cell carcinoma. Sci Transl Med, 2012. 4(137): p. 137ra75.

54. Contreras-Zarate, M.J., et al., Development of Novel Patient-Derived Xenografts from Breast Cancer Brain Metastases. Front Oncol, 2017. 7: p. 252.

55. Ni, J., et al., Combination inhibition of PI3K and mTORC1 yields durable remissions in mice bearing orthotopic patient-derived xenografts of HER2-positive breast cancer brain metastases. Nat Med, 2016. 22(7): p. 723–6.

56. Oshi, M., et al., Novel Breast Cancer Brain Metastasis Patient-Derived Orthotopic Xenograft Model for Preclinical Studies. Cancers (Basel), 2020. 12(2).

57. Morton, C.L. and P.J. Houghton, Establishment of human tumor xenografts in immunodeficient mice. Nat Protoc, 2007. 2(2): p. 247–50.

58. Nolte, S.M., et al., A cancer stem cell model for studying brain metastases from primary lung cancer. J Natl Cancer Inst, 2013. 105(8): p. 551–62.

59. Krepler, C., et al., A Comprehensive Patient-Derived Xenograft Collection Representing the Heterogeneity of Melanoma. Cell Rep, 2017. 21(7): p. 1953–1967.

60. McFadden, D.G., et al., Genetic and clonal dissection of murine small cell lung carcinoma progression by genome sequencing. Cell, 2014. 156(6): p. 1298–1311.

61. Cleary, A.S., et al., Tumour cell heterogeneity maintained by cooperating subclones in Wnt-driven mammary cancers. Nature, 2014. 508(7494): p. 113–7.

62. Sanborn, J.Z., et al., Phylogenetic analyses of melanoma reveal complex patterns of metastatic dissemination. Proc Natl Acad Sci U S A, 2015. 112(35): p. 10995–1000.

63. Marjanovic, N.D., et al., Emergence of a High-Plasticity Cell State during Lung Cancer Evolution. Cancer Cell, 2020. 38(2): p. 229–246 e13.

64. Gundem, G., et al., The evolutionary history of lethal metastatic prostate cancer. Nature, 2015. 520(7547): p. 353–357.

65. Faria, C.C., et al., Identification of alsterpaullone as a novel small molecule inhibitor to target group 3 medulloblastoma. Oncotarget, 2015. 6(25): p. 21718–29.

66. Silva, A., et al., Regulation of PTEN by CK2 and Notch1 in primary T-cell acute lymphoblastic leukemia: rationale for combined use of CK2- and gamma-secretase inhibitors. Haematologica, 2010. 95(4): p. 674–8.

67. Schindelin, J., et al., Fiji: an open-source platform for biological-image analysis. Nat Methods, 2012. 9(7): p. 676–82.

