## Supplementary Material for "Patient-derived models of brain metastases recapitulate the histopathology and biology of human metastatic cancers"

### SUPPLEMENTARY TABLES AND FIGURE LEGENDS

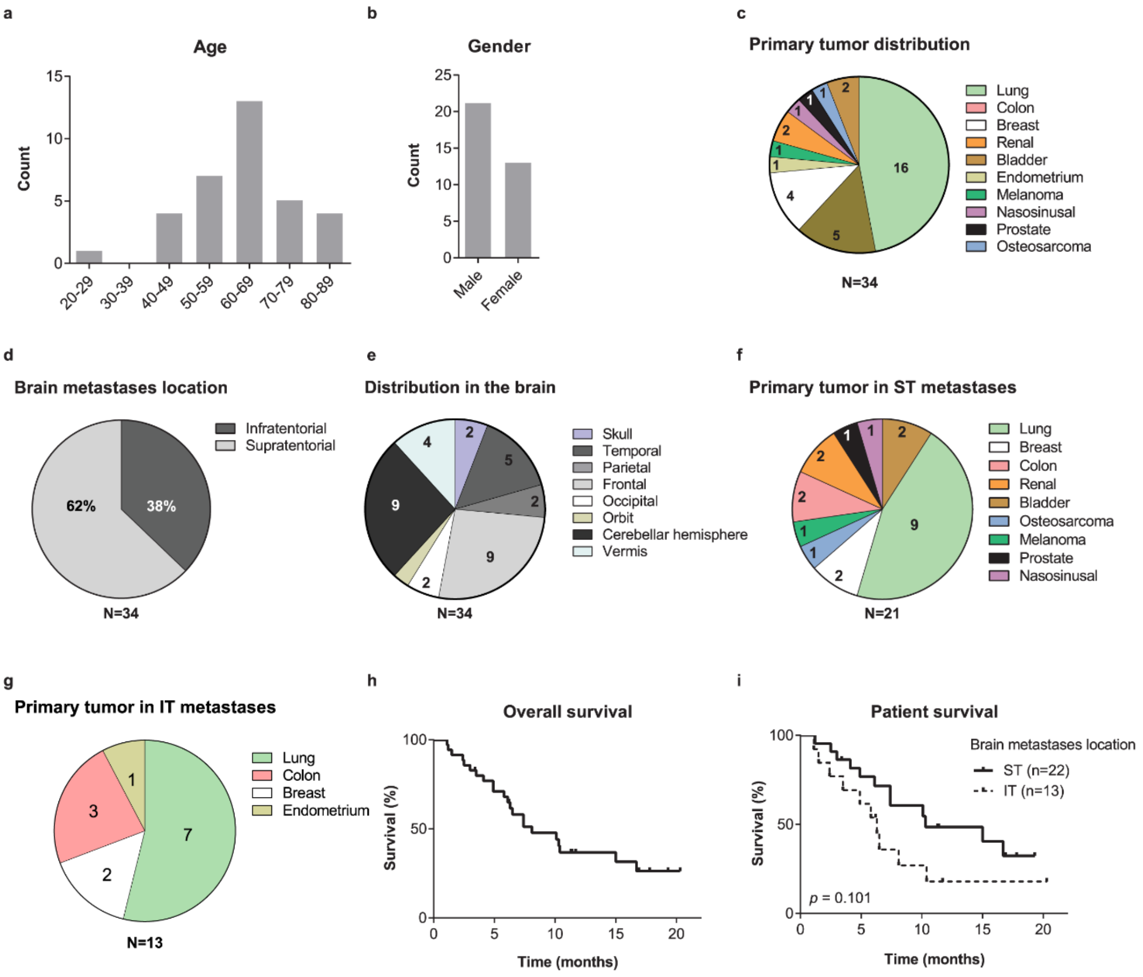

**Supplementary Figure 1.** Clinical characterization of the cohort of patients with BMs. (**A**) Age of patients at the time of BMs surgery. (**B**) Gender distribution. (**C**) Distribution of patients according to the primary tumor. (**D**) Percentage of BMs in the supratentorial and infratentorial compartments. (**E**) Anatomical location of BMs. (**F-G**) Distribution of primary tumors according to the location in the intracranial compartments. (**H**) Overall survival of patients upon diagnosis of BMs. (**I**) Patient overall survival according to the metastases location in the intracranial compartments. Differences were considered statistically significant for p-values<0.05, according to the Log-rank (Mantel-Cox) test.

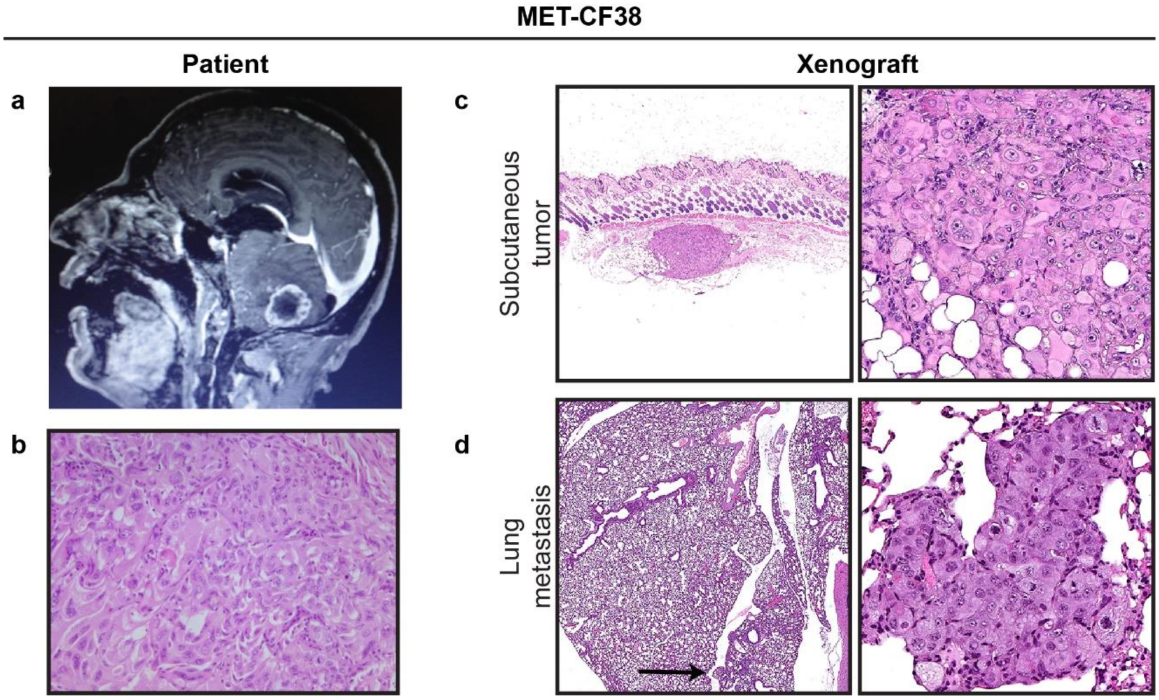


**Supplementary Figure 2.** Representative clinical case of a 49 years-old male patient with a stage IV lung adenocarcinoma where the BM xenografted in the flank originated spontaneous metastases in the mice lungs.

(**A**) Brain MRI, sagittal T1 contrast-enhanced sequence, showing a BM in the cerebellum. (**B**) H&E staining of the patient BMs. (**C**) Corresponding H&E staining of the mouse flank xenograft. (**D**) Exclusive dissemination to the mouse lungs through passaging.

**Supplementary Table 1.** Matched patient and PDX metastatic sites in subcutaneous PDX models.

| Primary tumor | Sample ID | Metastases location | | | | | | | | | |
|---|---|---|---|---|---|---|---|---|---|---|---|
| | | CNS | Lungs | Liver | Spleen | Kidney | Intraperit. | Lymph nodes | Soft tissues | Gonads | Bone/ Bone marrow |
| Lung | MET-CF38 | Green | Grey | | | | | | | | |
| | MET-CF58 | Green | Grey | Green | Green | | | | | | |
| | MET-CF66 | Green/Grey | Grey | Grey | Grey | Grey | | Green/Grey | | | Grey |
| | MET-CF68 | Green | Grey | | | Grey | Grey | Green/Grey | | | |
| | MET-CF78 | Green/Grey | Grey | Green/Grey | Grey | Grey | Green/Grey | Green/Grey | Grey | Grey | |
| | MET-CF80 | Green | | Grey | | Grey | | | Green/Grey | | |
| | MET-CF85 | Green | Grey | Grey | Grey | Grey | | | | | |
| | MET-CF87 | Green | | | | | | | Grey | | |
| | MET-CF90 | Green/Grey | Grey | Grey | Grey | Grey | | | | | |
| Melanoma | MET-CF69 | Green | Green/Grey | Green | | Grey | | | Grey | | |
| Colon | MET-CF70 | Green | Green | | | | | | Grey | | |
| | MET-CF82 | Green | Green | Green | | | | Green | Grey | | |
| Endometrium | MET-CF81 | Green | Green/Grey | | | | Grey | | | | |
| Breast | MET-CF84 | Green | Grey | | | | | | | | |

Green - organs with tumor involvement in the patient; Grey - organs with tumor involvement in the mouse.

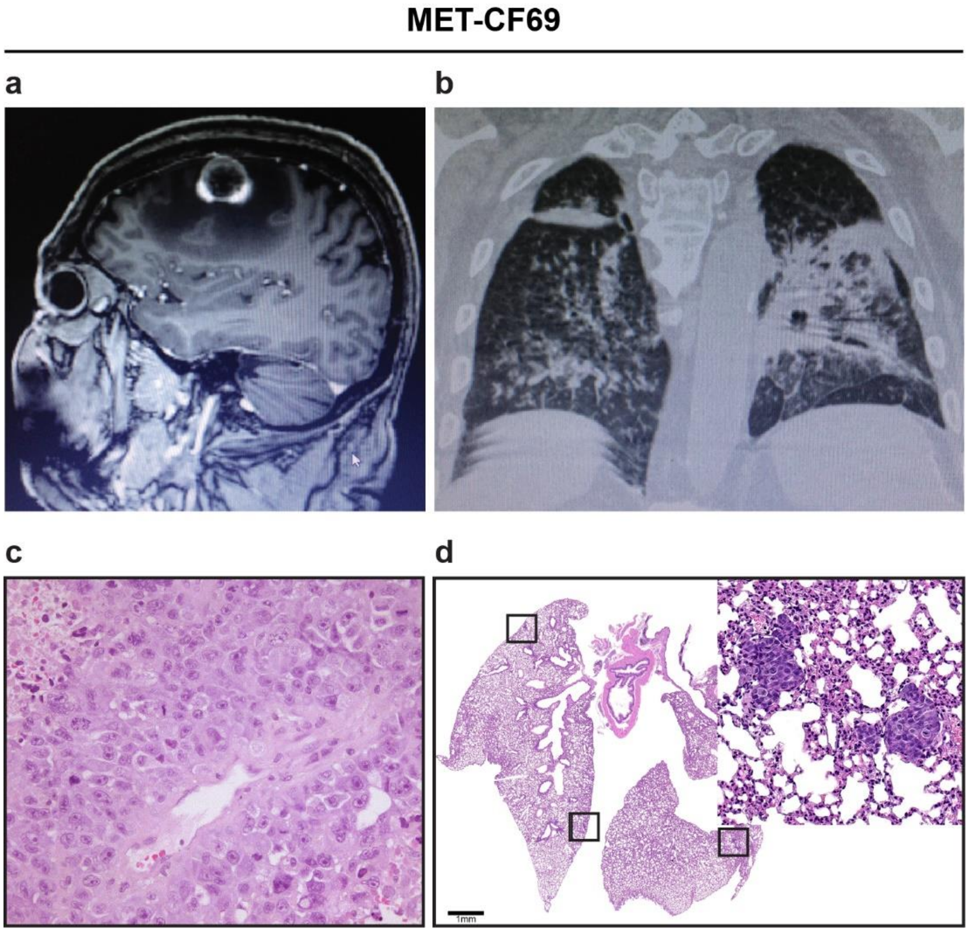


**Supplementary Figure 3.** Representative clinical case of a 76 years-old male patient with metastatic melanoma to the brain and to the lungs, where the xenografted BM originated spontaneous metastases in the mice lungs. (**A**) Magnetic Resonance Imaging (MRI) of the brain, sagittal T1 contrast-enhanced, showing a left frontal BM. (**B**) Computed Tomography (CT) scan of the patient's lungs showing a diffuse infiltration by cancer cells. (**C**) H&E staining of the patient BMs (20x). (**D**) H&E staining of the mice lungs exhibiting multiple metastases.

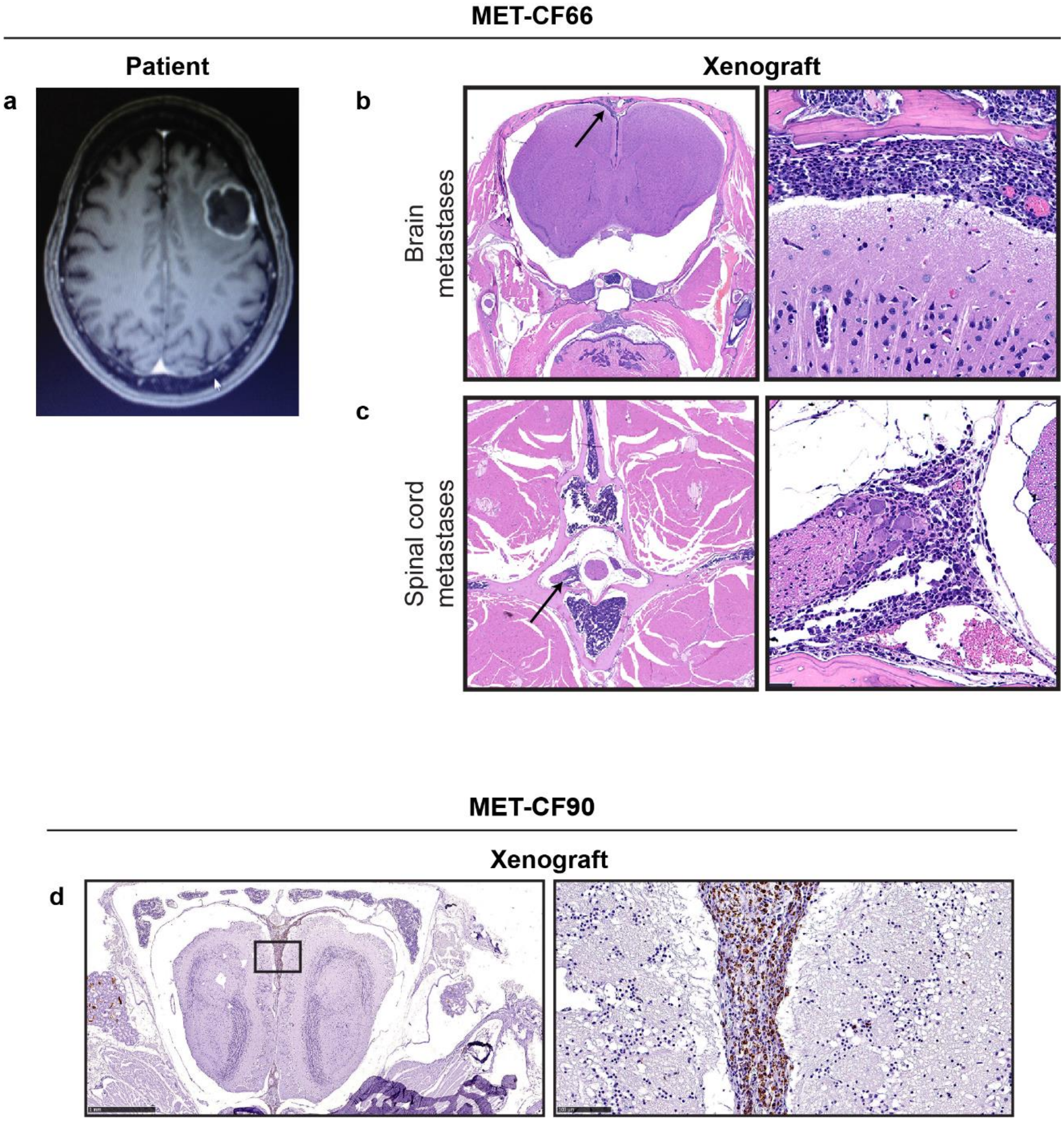


**Supplementary Figure 4.** Spontaneous leptomeningeal dissemination in flank implanted BMs from lung cancer.

(**A**) Brain MRI, axial T1 contrast-enhanced sequence, of a 66 years-old female patient with a left frontal BM from lung adenocarcinoma. (**B-C**) H&E staining showing spontaneous leptomeningeal dissemination to the brain and the spinal cord in the matched xenograft. (**D**) Representative anti-human mitochondria staining of a mouse brain with leptomeningeal dissemination derived from a 68 years-old patient with a posterior fossa BM from lung adenocarcinoma.

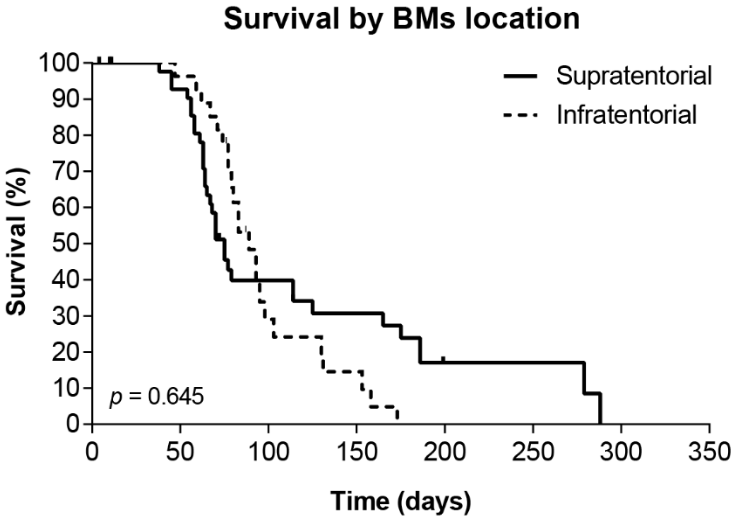


**Supplementary Figure 5.** Overall survival of intracardiac PDXs according to the location of patient's metastases in the supratentorial and infratentorial compartments. Differences were considered statistically significant for p-values<0.05, according to the Log-rank (Mantel-Cox) test.

**Supplementary Table 2.** Matched patient and PDX metastatic sites in intracardiac PDX models.

| Primary tumor | Sample ID | Metastases location | | | | | | | | | | | | |
|---|---|---|---|---|---|---|---|---|---|---|---|---|---|---|
| | | CNS | Lungs | Liver | Kidney | Suprarenal | Intraperit. | Spleen | Gonads | Orbita | Mandibula | Soft tissues | Bone/Bone marrow | Subcut. |
| Lung | MET-CF30 | Green | | Grey | | Green | Green | | | | | | | |
| | MET-CF66 | Green | | | | | Grey | Grey | | | | | | |
| | MET-CF68 | Green | Grey | Grey | Grey | Grey | | | | | | Grey | Grey | |
| | MET-CF78 | Green | Grey | Green | Grey | | Green/Grey | | | | | Grey | | Grey |
| | MET-CF79 | Green | | | | | | | | | | | | Grey |
| | MET-CF87 | Green/Grey | | Grey | | | | | | | | | | |
| Osteosarcoma | MET-CF65 | Green | Green/Grey | Grey | Grey | Grey | | | | | | | | |
| Melanoma | MET-CF69 | Green/Grey | Green/Grey | Green/Grey | Grey | Grey | Grey | | | Grey | | Grey | Grey | |
| | MET-CF70 | Green/Grey | Green/Grey | Grey | Grey | Grey | Grey | | Grey | | Grey | Grey | Grey | |
| Colon | MET-CF82 | Green | Green | Green | | Grey | | | | | | | Grey | |
| | MET-CF89 | Green/Grey | Green | | Grey | Green/Grey | | | Grey | | | | Grey | |
| Bladder | MET-CF29 | Green/Grey | | | | | | | | | | | | |
| Endometrium | MET-CF81 | Green/Grey | Green/Grey | Grey | Grey | Green | Grey | | Grey | | | Grey | Grey | |

Green - organs with tumor involvement in the patient; Grey - organs with tumor involvement in the mouse.

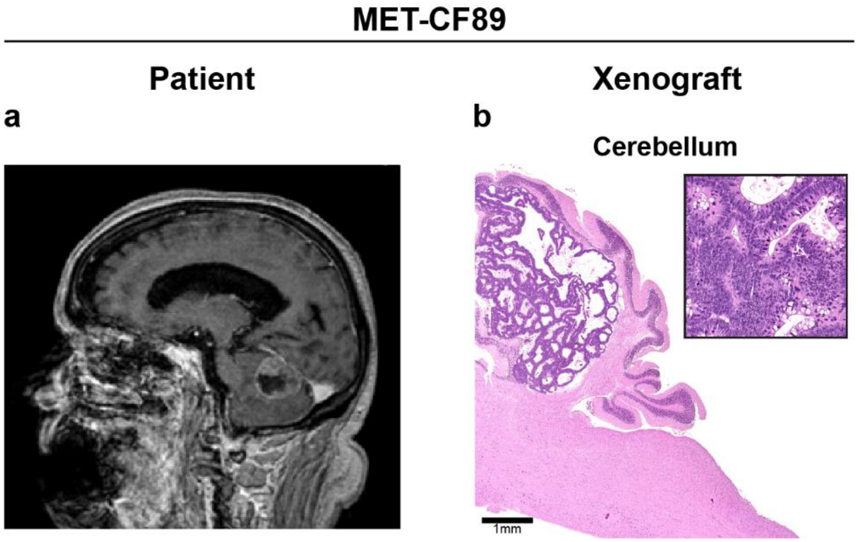


**Supplementary Figure 6.** Intracardiac xenograft of a surgically resected cerebellar BM from a patient with colon cancer mirrored the intracranial location of the patient's tumor. (**A**) Brain MRI, sagittal T1 contrast-enhanced sequence, of a 81 years-old male patient with a BM in the cerebellum from a colon carcinoma. (**B**) Corresponding H&E staining of a metastasis in the mouse cerebellum.

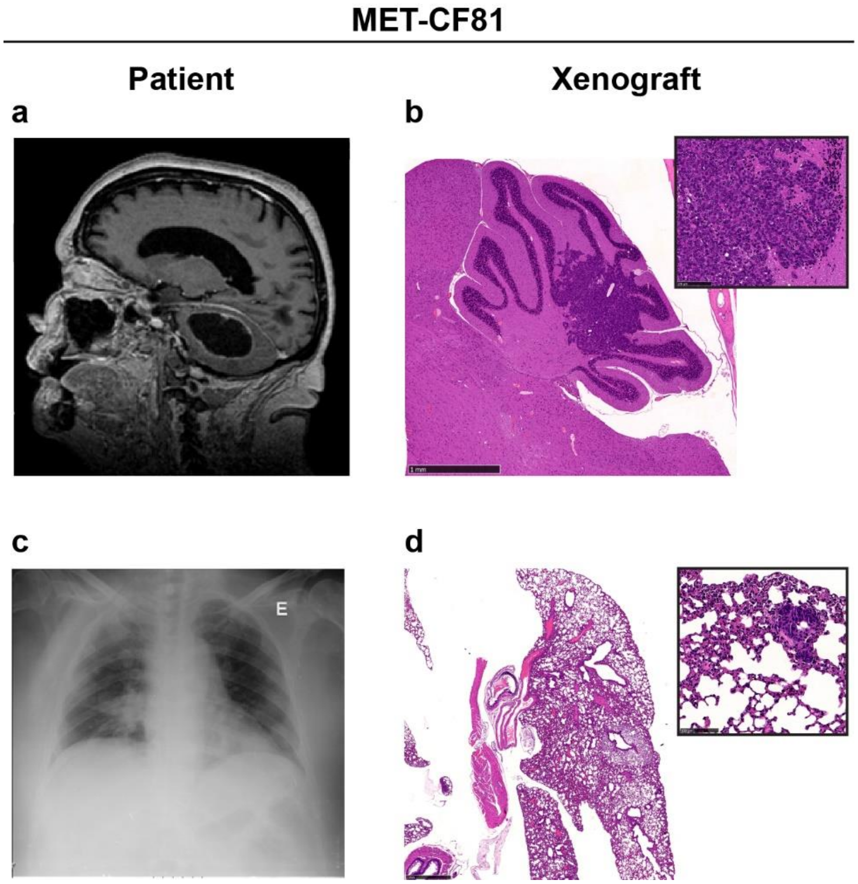


**Supplementary Figure 7.** Representative clinical case of a 67 years-old female patient with metastatic endometrium cancer, where the intracardiac xenograft shared two metastatic sites with the donor. (**A**) Brain MRI, sagittal T1 contrast-enhanced sequence, showing a cystic BM in the cerebellum and (**B**) the correspondent H&E staining of the mouse metastasis in the cerebellum. (**C**) Right lung metastases in the patients' chest X Ray and (**D**) the matched lung metastases in the xenograft.
